# Inhibiting endoplasmic reticulum stress decreases tumor burden in a mouse model for hepatocellular carcinoma

**DOI:** 10.1101/826453

**Authors:** Natasa Pavlovic, Carlemi Calitz, Kessarin Thanapirom, Giuseppe Mazza, Krista Rombouts, Pär Gerwins, Femke Heindryckx

## Abstract

Hepatocellular carcinoma (HCC) is a liver tumor that arises in patients with cirrhosis. Hepatic stellate cells are key players in the progression of HCC, as they create a fibrotic micro-environment and produce growth factors and cytokines that enhance tumor cell proliferation and migration. We assessed the role of endoplasmic reticulum (ER) stress in the cross-talk between stellate cells and HCC-cells. Mice with a fibrotic HCC were treated with the IRE1α-inhibitor 4μ8C, which reduced tumor burden and collagen deposition. By co-culturing HCC-cells with stellate cells, we found that HCC-cells induce ER-stress in stellate cells, thereby contributing to their activation. Inhibiting IRE1α blocked stellate cell activation, which inhibited tumor cell proliferation and migration in different *in vitro* 2D and 3D co-cultures. Our results suggest that IRE1α is an important mediator in the communication between stellate cells and cancer cells and components of the ER-stress pathway may be therapeutically relevant for HCC-patients.

**Impact statement:** IRE1α is an important mediator in the communication between stellate cells and cancer cells and components of the ER-stress pathway may be therapeutically relevant for liver cancer.

## INTRODUCTION

Hepatocellular carcinoma (HCC) is a primary liver tumor that typically arises in a background of chronic liver disease and cirrhosis (1). One of the key players in the progression of cirrhosis to HCC is the hepatic stellate cell, which activates during liver damage and differentiates towards a contractile myofibroblast-like cell responsible for the deposition of extracellular matrix proteins (ECM) such as collagen (2). Activated stellate cells can induce phenotypic changes in cancer cells through the production of growth factors and cytokines that stimulate tumor cell proliferation and induce a pro-metastatic phenotype (3). One of the key factors in the cross talk between tumor cells and stellate cells is tumor growth factor beta (TGFβ) (4-6). Malignant hepatocytes secrete high levels of TGFβ, which can contribute to the activation of stellate cells in the nearby stroma. These activated stellate cells are then responsible for the deposition of ECM. Several of the ECM components such as proteoglycans, collagens, laminin, and fibronectin interact with tumor cells and cells in the stroma, which can directly promote cellular transformation and metastasis (7, 8). The ECM can also act as a reservoir for growth factors and cytokines, which can be rapidly released to support the tumor’s needs. In addition, activated stellate cells contribute to a highly vascularized tumor micro-environment, by secreting pro-angiogenic molecules and by recruiting pro-angiogenic (and pro-tumoral) myeloid and lymphoid derived cell types (9). By constricting the hepatic microvasculature, they also cause hypoxia, which contributes to the angiogenic switch and can induce a more aggressive tumor phenotype (10). It is therefore not surprising that tumor cells actively secrete growth factors (such as TGFβ) to induce activation and migration of stellate cells, which creates a fibrotic environment that further supports and enhances tumor progression (2, 11, 12). Since activated stellate cells play an essential role in the onset and progression of HCC, blocking their activation has been proposed as a potential therapy for patients with HCC (13). One strategy to block stellate cell activation, is by targeting the unfolded protein response (UPR).

The unfolded protein response serves to cope with misfolded or unfolded proteins in the ER in an attempt to restore protein folding, increase ER-biosynthetic machinery and maintain cellular homeostasis (14). It can exert a cytoprotective effect by re-establishing cellular homeostasis, while apoptotic signaling pathways will be activated in case of severe and/or prolonged ER-stress (15). The presence of misfolded proteins is sensed via 3 transmembrane proteins in the ER: IRE1α, PERK and ATF6a. Actors of the ER-stress pathways have been described to play a role in the progression of solid tumors, such as breast cancer (16), colon cancer (17) and HCC (18). Activation of the UPR has also been shown to affect different fibrotic diseases (19), including non-alcoholic fatty liver disease (20-22), hepatitis B-induced carcinogenesis (23) and biliary cirrhosis (24). We have previously shown that inhibiting the IRE1α-branch of the UPR-pathway using 4μ8C, blocks TGFβ-induced activation of fibroblasts and stellate cells *in vitro* and reduces liver fibrosis *in vivo* (25). In the current study, our aim was to define the role of ER-stress in the cross-talk between hepatic stellate cells and tumor cells in liver cancer. We show that pharmacologic inhibition of the IRE1α signaling pathway decreases tumor burden in a chemically induced mouse model for HCC. Using several *in vitro* co-culturing methods, we identified that tumor cells induce ER-stress in hepatic stellate cells. Blocking ER-stress in these hepatic stellate cells prevents their activation and decreases proliferation and migration of tumor cells co-cultured with hepatic stellate cells.

## MATERIAL AND METHODS

### Mouse model

A chemically induced mouse model for HCC was used, as previously described (26, 27). Briefly, 5-week-old male sv129 mice received intraperitoneal injections once per week with 35mg/kg bodyweight DEN diluted in saline. From week 10, mice were injected twice per week with 10μg/g bodyweight 4μ8C (Sigma) in saline. After 25 weeks mice were euthanized and samples were taken for analysis. This method was approved by the Uppsala ethical committee for animal experimentation (C95/14). Each group contained 8 mice, which generates enough power to pick up statistically significant differences between treatments, as determined from previous experience (26, 27). Mice were assigned to random groups before treatment.

### Olink multiplex proximity extension assay

Liver samples were homogenized in ice-cold RIPA containing protease inhibitors (Sigma Aldrich). Homogenates were kept on ice for 20–30 min, whilst mixing vigorously to enhance disruption of the cell membranes. The homogenates were centrifuged (20 min, 13 000 rpm, 4°C) and supernatant containing protein was collected. Supernatant was stored at -20°C until protein measurement. Protein concentration was measured using the BCA kit (ThermoFisher) and all samples were diluted to 1 mg/mL protein in RIPA. Samples from 3 biological replicates per group were analyzed with a multiplex proximity extension assay for ninety-two biomarkers in the murine exploratory panel (Olink Bioscience, Uppsala, Sweden) (28). Samples were loaded random on the assay plates. Raw data was deposited in Dryad (29).

### Cell culture and reagents

The HCC-cell lines (HepG2, ATCC and Huh7, kind gift from Dilruba Ahmed, Karolinska Institute) and hepatic stellate cell-line LX2 (Sigma-Aldrich, Darmstadt, Germany) were cultured at 37°C with 5% CO_2_ in Dulbecco modified eagle medium (DMEM) supplemented with 10% fetal bovine serum (FBS) (ThermoFisher, Stockholm, Sweden). No FBS was used during starvation and stimulation with growth factors. Cells were detached using trypsin-EDTA (ThermoFisher), re-suspended in growth medium and plated at a density of 5×10^3^ cells/cm^2^. Cells were allowed to attach and left undisturbed for 8h before being starved for 16h. Afterwards, fresh starvation medium containing indicated growth factors or substances were added. Cells were exposed for 48h to 100μM 4μ8C (Sigma-Aldrich) or 10μM SB-431541 (Tocris, Abingdon, UK) as previously described (25). For transwell co-culturing experiments, cells were grown on 12-well Corning Tissue Culture-plates with transwell inserts (Sigma-Aldrich) with 0,4μm-pore size, allowing the exchange of soluble factors, but preventing direct cell contact.

3D-tumor spheroids were generated on 96-well ultra-low attachment plates (Sigma-Aldrich) (30). After 6 days, spheroids had reached approximately 1mm^2^ and 4μ8C was added. Proliferation was monitored during the subsequent 4 days. Tumor spheroids were retrieved from the plates after 10 days.

Fluorescent labeling of cells was done in imaging experiments by using CellTracker (ThermoFisher), according to manufacturer’s instructions. Cell pellets were incubated 30 minutes with 1μM of CellTracker™ Red CMTPX or 1μM of CellTracker™ Green CMFDA. Cells were washed twice in PBS prior to co-culturing.

### Human liver scaffold decellularization and cell culture usage

Human healthy livers were obtained under the UCL Royal Free BioBank Ethical Review Committee (NRES Rec Reference: 11/WA/0077) approval. Informed consent was obtained for each donor and confirmed via the NHSBT ODT organ retrieval pathway (31). Liver 3D-scaffolds, were decellularized, sterilized and prepared for cell culture use as preciously described (31). LX2 and HepG2-cells, as either mono-cultures or mixed co-culture, were released on top of each scaffold as 2.5*10^5^ cells in 20μL (32).

### Proliferation

Cell proliferation was monitored via a resazurin reduction assay. A 1%-resazurin solution was added in 1/80 dilution to the cells and incubated for 24h, after which fluorescent signal was measured with a 540/35 excitation filter and a 590/20 emission filter on a Fluostar Omega plate reader.

### Transfections

Nucleofection with 0,1μM siIRE1α (s200432, ThermoFisher), or 0,1 μM siCtrl (4390843, ThermoFisher) was done using Amaxa Nucleofector program S-005 in Ingenio electroporation solution (Mirus Bio LLC, Taastrup, Denmark).

### Migration and chemotaxis

Non-directional migration was assessed using a scratch wound assay on fluorescently labelled LX2-cells and HepG2-cells. Scratch size was measured by analyzing light microscopy images in ImageJ, using the MRI Wound Healing Tool plug-in (http://dev.mri.cnrs.fr/projects/imagej-macros/wiki/Wound_Healing_Tool). Image analysis was done in ImageJ.

Directional migration was assessed using CellDirector-devices (GradienTech, Uppsala, Sweden). HepG2 and LX2-cells were labelled with CellTracker-dye and left to adhere overnight in the CellDirector-devices. Non-adherent cells were washed away with DMEM and cells were starved for 1h prior to commencing experiments. A gradient of 0 to 10% FBS was created with a flow rate of 1.5 µl/minute. Cell movement was recorded using an Axiovision 200M microscope (Zeiss, Stockholm, Sweden) for 4h and tracked using Axiovision software (Zeiss). During the assay cells were kept at 37°C with 5% CO_2_.

### Quantitative RT-PCR of mRNA

RNA was isolated from tissue or cell culture using the EZNA RNA isolation Kit (VWR, Spånga, Sweden) or using TRIzol reagent and RNeasy Universal Mini Kit (Qiagen, Sollentuna, Sweden) for human liver scaffolds (31). RNA-concentration and purity were evaluated using Nanodrop. Afterwards, 500ng of mRNA was reverse transcribed using iScript cDNA synthesis kit (Bio-rad, Solna, Sweden). Amplifications were done using primers summarized in supplementary table 1. mRNA-expression was normalized to *18S, GAPDH* and/or *TBP1*. Fold change was calculated via the delta-delta-CT method, by using the average CT value of 3 technical replicates.

The procedure to detect the spliced and unspliced isoforms of XBP1 was done by digesting RT-PCR product with the restriction enzyme *Pst-*I (ThermoFisher). This cleaves unspliced-XBP1 containing the Pst-I-cleavage site (CTGCA^G), but leaves the spliced isoform intact. The digestion reaction was stopped after 18h by 0,5M EDTA (pH 8.0) and run on a 1,5% agarose gel for 1h at 180V.

### Stainings and immunocytochemistry

Tissue samples were fixed in 4% paraformaldehyde for 24h and subsequently embedded in paraffin. Cells and tumor spheroids were fixed for 10 minutes in 4% paraformaldehyde and stored at 4°C. Paraffin embedded tissue samples were cut at 5μm and dried overnight. Sections were de-paraffinized and rehydrated prior to staining. Collagen was stained using the picrosirius red staining with an incubation time of 30 minutes, followed by 10 minutes washing in distilled water. Haematoxilin-eosin (H&E) staining was done according to standard practice. Images were acquired using a Nikon eclipse 90i microscope equipped with a DS-Qi1Mc camera and Nikon plan Apo objectives. NIS-Elements AR 3.2 software was used to save and export images. Quantification of collagen deposition was performed blindly with ImageJ software by conversion to binary images after color de-convolution to separate Sirius Red staining, as previously described (33).

Paraformaldehyde fixed cells and spheroids were washed with tris-buffer saline (TBS) and blocked for 30 minutes using 1% bovine serum albumin in TBS + 0,1% Tween. For liver tissue, antigen retrieval was done at 95°C in sodium citrate buffer and endogenous mouse IgG was blocked using a rodent blocking buffer (ab127055, abcam) following manufacturer’s guidelines. Blocking was followed by an overnight incubation at 4°C with antibodies against α-smooth muscle actin (αSMA) (clone 1A4, Sigma), Bip (ab21685, abcam) or p-IRE1α (PAB12435, Abnova). A 40-minute incubation was used for the secondary antibody (Rabbit anti-mouse Alexa Fluor-488 or donkey anti-rabbit Alexa Fluor-633) and cell nuclei were stained with Hoechst for 8 minutes. Images were taken using an inverted confocal microscope (LSM 700, Zeiss) using Plan-Apochromat 20× objectives and the Zen 2009 software (Zeiss). The different channels of immunofluorescent images were merged using ImageJ software. Quantifications were done blindly with ImageJ software by conversion to binary images for each channel and automated detection of staining on thresholded images using a macro.

For histological and immunohistochemical analysis of the human liver scaffolds, 4μm slides were cut from paraffin embedded blocks. The sections were de-paraffinized and rehydrated prior to staining. To retrieve the antigens, slides were microwaved at high power for 5 minutes in pre-heated 10 mM sodium citrate buffer, and subsequently left to cool down to room temperature. Following this, a single wash was performed in 100 mM Glycine in PBS, after which the slides were blocked for 2h in TNB Blocking Reagent (Ancillary Products, FP1020). Slides were then incubated for 2h in the following antibodies; Ki67 (1:100; eBioscience™, SolA15), and EPCAM (1:100; Abcam, ab71916). A 1h incubation was used for the secondary antibody (goat anti-rat Alexa Fluor 555 and Rabbit anti-mouse Alexa Fluor 488). Sections were mounted with Fluoromount-G™, with DAPI (Invitrogen, 00-4959-52). Images were taken with using an inverted confocal microscope (LSM 780, Zeiss) using Plan-Apochromat 10× objectives and the Zen 2009 software (Zeiss).

### Enzyme-Linked immune Sorbent Assay (ELISA)

Medium samples from cells and from the engrafted scaffolds were used to measure TGFβ using ELISA (88-8350-22, ThermoFisher), following manufacturer’s guidelines. The average from 2 technical replicates were used for calculations.

### SDS-PAGE and western blot

Protein lysates in lysis buffer were mixed with 2x laemmli buffer and heated to 95°C for 5 minutes before being loaded onto a 10% polyacrylamide gel. After separation, the proteins were transferred to an Immobilon-Fl membrane (Millipore). The membrane was blocked using the Odyssey blocking buffer (Licor) diluted 1:4 in PBS, and then incubated with primary and secondary antibodies. After primary and secondary antibody incubation the membrane was washed 3×15 minutes in PBS-T (Phosphate buffered saline (Gibco), 0.1% Tween-20). Primary antibodies used were Bip (ab21685, abcam), p-IRE1α (PAB12435, Abnova) or vinculin (14-9777-82, ThermoFisher) all added in blocking buffer with 0.1% Tween-20. Secondary antibodies used were: goat-anti-mouse alexa 680 (Invitrogen) and goat-anti-rabbit IRDye 800 (Rockland) 1:20 000 diluted in blocking buffer with 0.1% Tween-20 and 0.01% SDS. All incubations were carried out at room temperature for 1h or overnight at 4°C. The membranes were scanned using an Odyssey scanner (LI-COR Biotechnology) and band intensities quantified using the Odyssey 2.1 software and normalized to the vinculin signal in each sample.

### Gene-set enrichment analysis

Gene expression profiles of HCC with a fibrous stroma and without fibrous stroma was accessed through PubMed’s Gene Expression Omnibus via accession number GSE31370 (34). A gene-set containing 79 proteins involved in the unfolded protein response was downloaded from The Harmonizome (35) and GSEA software was used to perform a gene-set enrichment assay (36).

### Human protein atlas

Images from biopsies from HCC patients stained with antibodies against WIPI1 (37), SHC1 (38), PPP2R5B (39) and BiP (40) were obtained through the Human Protein Atlas (41).

### Statistics

Data are presented as mean ± s.e.m. Statistical significance was determined using an unpaired, two-tailed Student’s T-test or one-way analysis of variance (ANOVA) followed by Tukey’s multiple comparison test. Survival curves were generated with the Kaplan-Meier method and statistical comparisons were made using the log-rank method. P-values <0.05 were considered statistically significant. *In vitro* experiments were done in at least 3 biological replicates, which we define as parallel measurements of biologically distinct samples taken from independent experiments. Technical replicates we define as loading the same sample multiple times on the final assay. The *in vivo* experiments were done on at least 5 independent animals. Outliers were kept in the analyses, unless they were suspected to occur due to technical errors, in which case the experiment was repeated.

## RESULTS

### Pharmacological inhibition of IRE1α reduces tumor burden in a chemically induced mouse model for HCC

Hepatocellular carcinoma was induced in mice by weekly injections with diethylnitrosamine (DEN) for 25 weeks (26). From week 10, IRE1α-endonuclease activity was pharmacologically inhibited with 4μ8C. Histological analysis of liver tissue confirmed presence of liver tumors in a fibrotic background (Figure 1A). Treatment with 4μ8C significantly reduced tumor burden (Figure1B), as measured on H&E-stained liver slides (Figure1A). Stellate cell activation and liver fibrosis was quantified by Sirius Red staining (Figure 1A and 1C) and immunohistochemical staining with αSMA-antibodies (Figure 1A and 1D) on liver sections. Mice with HCC had a significant increase in the percentage of collagen (Figure 1C) and αSMA-staining (Figure 1D), compared to healthy mice. Treatment with 4μ8C restored collagen (Figure 1C) and αSMA-levels (Figure 1D and Figure 1E) to a similar level as healthy livers. mRNA expression levels of PCNA were determined on tumor nodules and surrounding non-tumor stromal tissue (Figure 1E). As expected, proliferation of cells was increased within the tumor itself, compared to the levels seen in healthy liver tissue and stromal tissue. Treatment with 4μ8C significantly decreased the levels of PCNA mRNA expression within the tumor, suggesting a decrease in tumor cell proliferation. A proteomics array using the Olink Mouse Exploratory assay revealed that DEN-induced murine tumors had a significantly increased protein expression of 20 oncogenic proteins compared to healthy controls (Figure 1F and table 1). In the 4μ8C-treated group, only 11 oncogenic proteins were increased compared to healthy controls (Figure 1F and table 1). Treatment with 4μ8C also significantly reduced the expression of two HCC promotors, Prdx5 and DDah1 (Figure 1F and table 1).

**Table 1:**
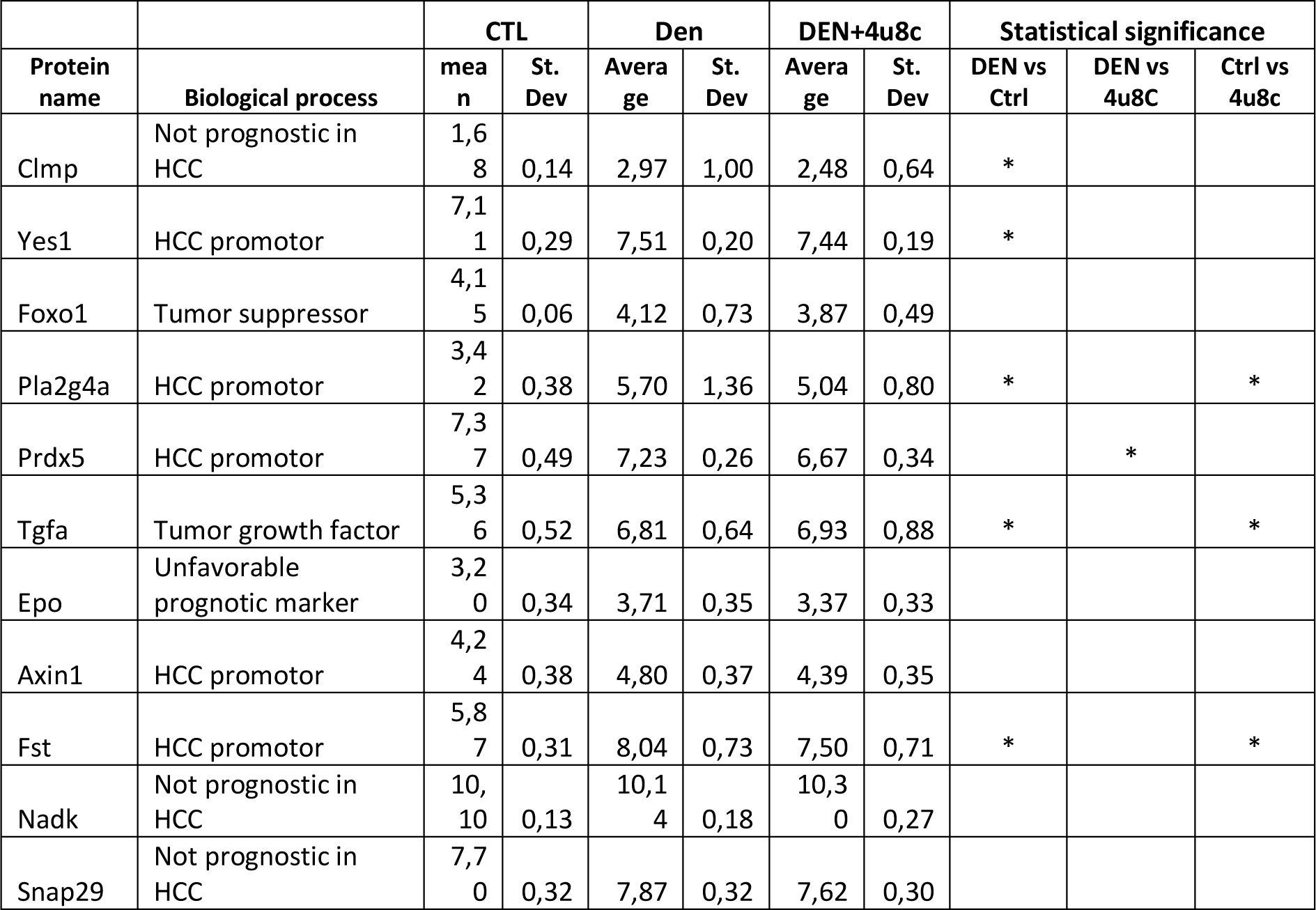
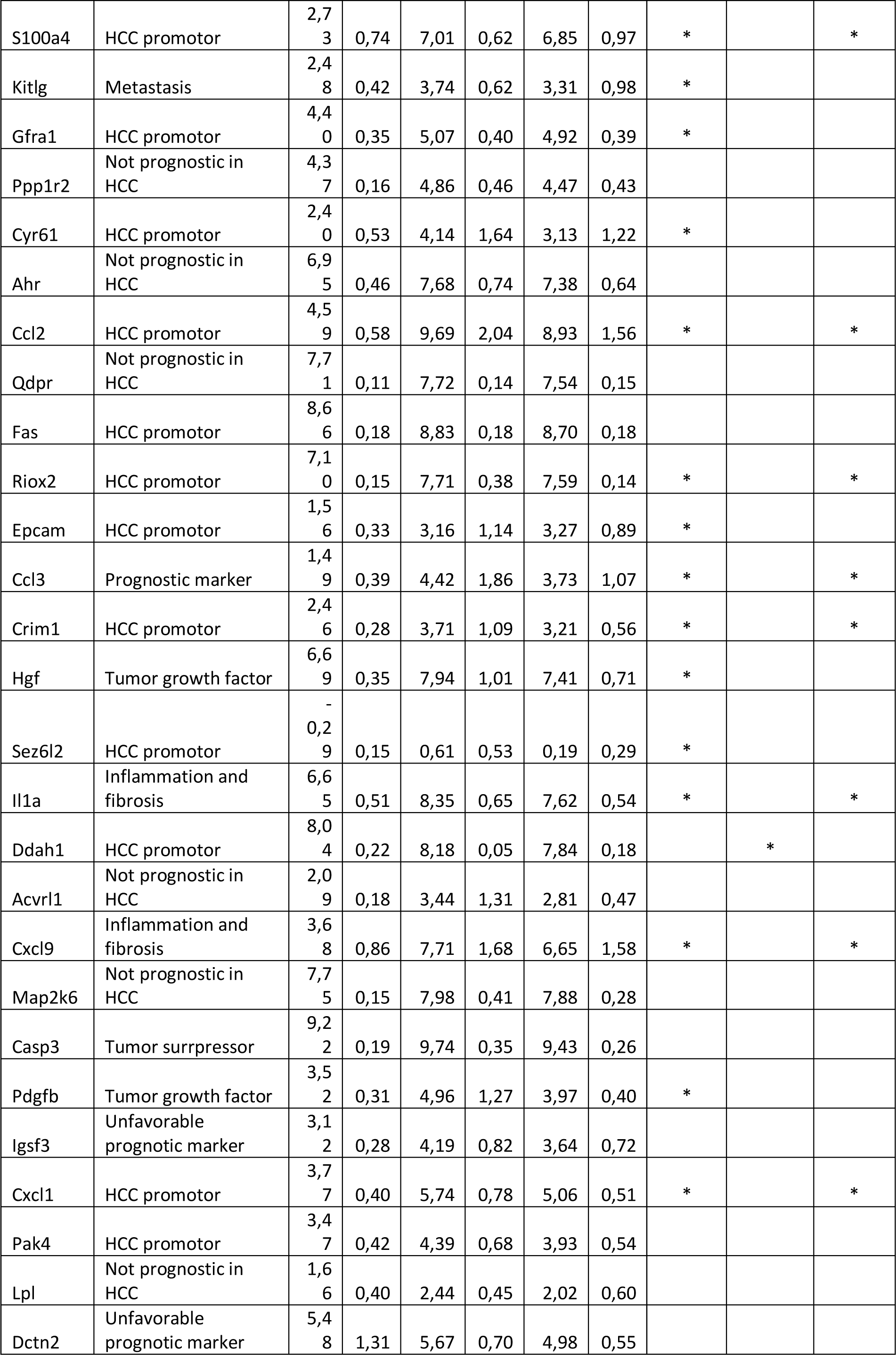
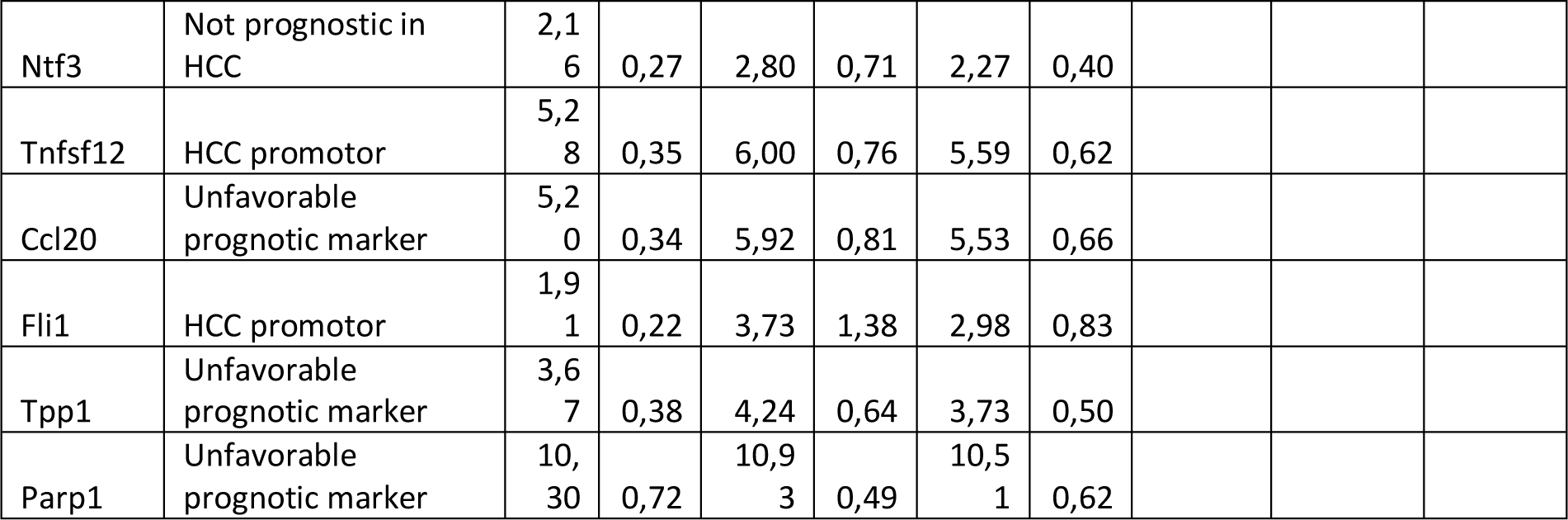
A proteomics array using the Olink Mouse Exploratory assay – source data figure 1F

**Fig. 1.**
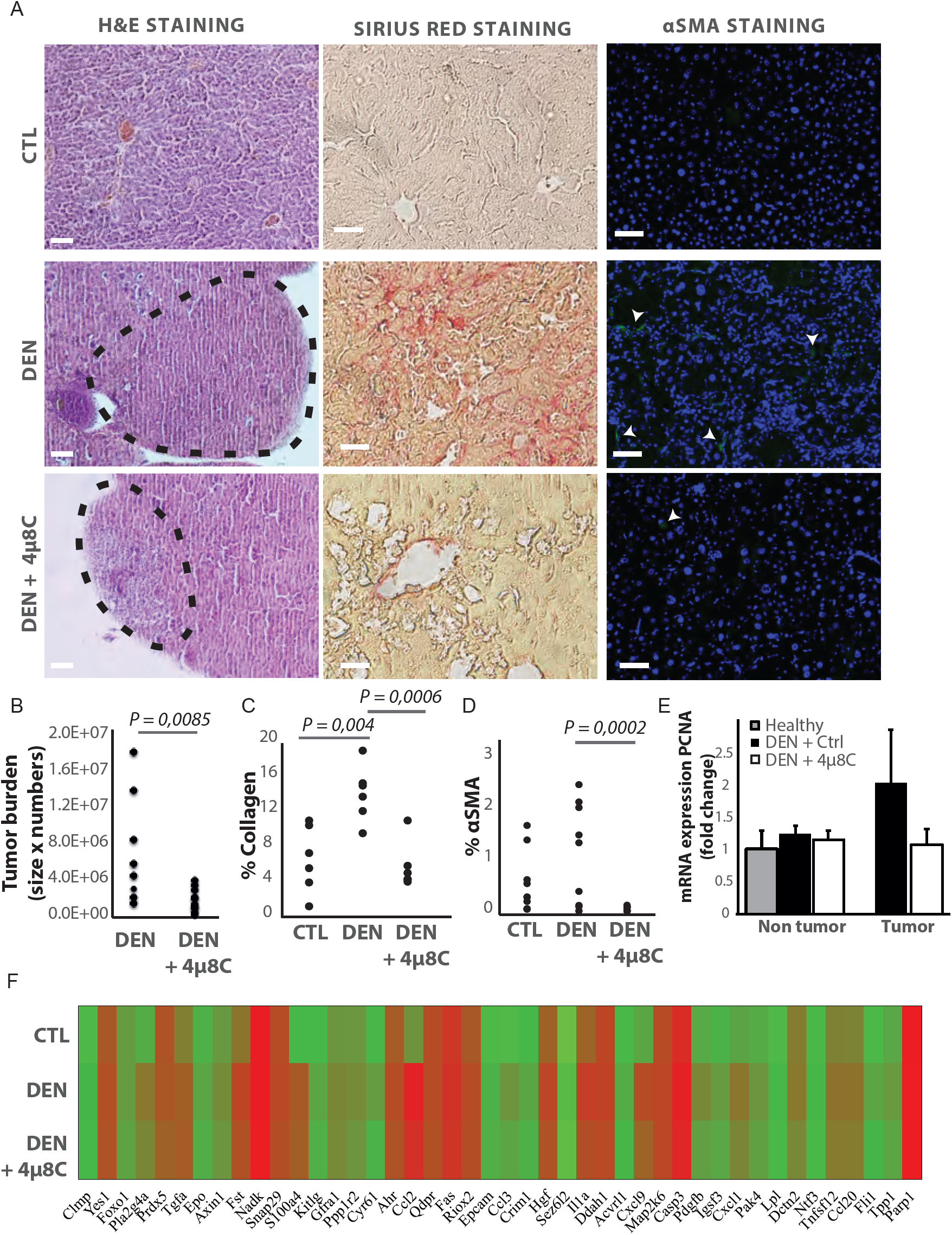
Inhibiting IRE1α reduces tumor burden *in vivo.* ***(A)*** Representative images of liver slides stained with hematoxylin and eosin (H&E), Sirius red and αSMA-antibodies. ***(B)*** tumor burden of mice with DEN-induced HCC treated with 4μ8C or vehicle-treated controls. ***(C)*** Quantification of percentage of collagen and ***(D)*** αSMA on liver slides. ***(E)*** mRNA expression of PCNA in liver tissue from mice with HCC treated with 4μ8C ***(F)*** Heatmap showing protein expression levels in healthy liver, DEN-induced HCC and DEN-induced HCC treated with 4μ8C from 3 biological replicates per group. P-values were calculated via the Student’s T-test, scale bars = 120μm.

### Markers of the unfolded protein response are upregulated in HCC and mainly located in the tumor stroma

mRNA-levels of the ER-stress-chaperone BiP were measured in tumor and surrounding non-tumor tissue of mice with DEN-induced HCC (Supplementary figure 1A). BiP-mRNA-expression was increased in the surrounding non-tumor tissue of DEN-induced mice, while there was no difference within the tumor, compared to healthy controls. Western blot confirmed the increase of BIP-protein expression in DEN-induced livers, which was reduced after treatment with 4μ8C (Supplementary figure 1 B). Co-staining of liver tissue with αSMA and p-IRE1α-antibodies (Supplementary figure 1C and D) or BiP-antibodies (Supplementary figure 1E) in untreated mice with HCC, revealed that expression of ER-stress markers was mainly localized within activated stellate cells in the liver.

A gene-set enrichment assay on microarray data from HCC-patients with fibrotic septae and without fibrotic septae showed an increase of genes involved in the UPR in the fibrotic HCC samples compared to non-fibrous HCC (supplementary figure 2A). Several actors of the IRE1α-branch of the UPR are amongst the genes that contribute to the core-enrichment of this analysis (table 2). Immunohistochemical staining of liver biopsies from HCC-patients further confirmed presence of IRE1α-mediated ER-stress markers BiP, PPP2R5B, SHC1 and WIPI1 localized in the fibrotic scar tissue and near hepatic blood vessels (Supplementary figure 2B). In addition, increased expression of these markers was significantly correlated with poor survival in patients with liver cancer (Supplementary figure 2C).

**Table 2:**
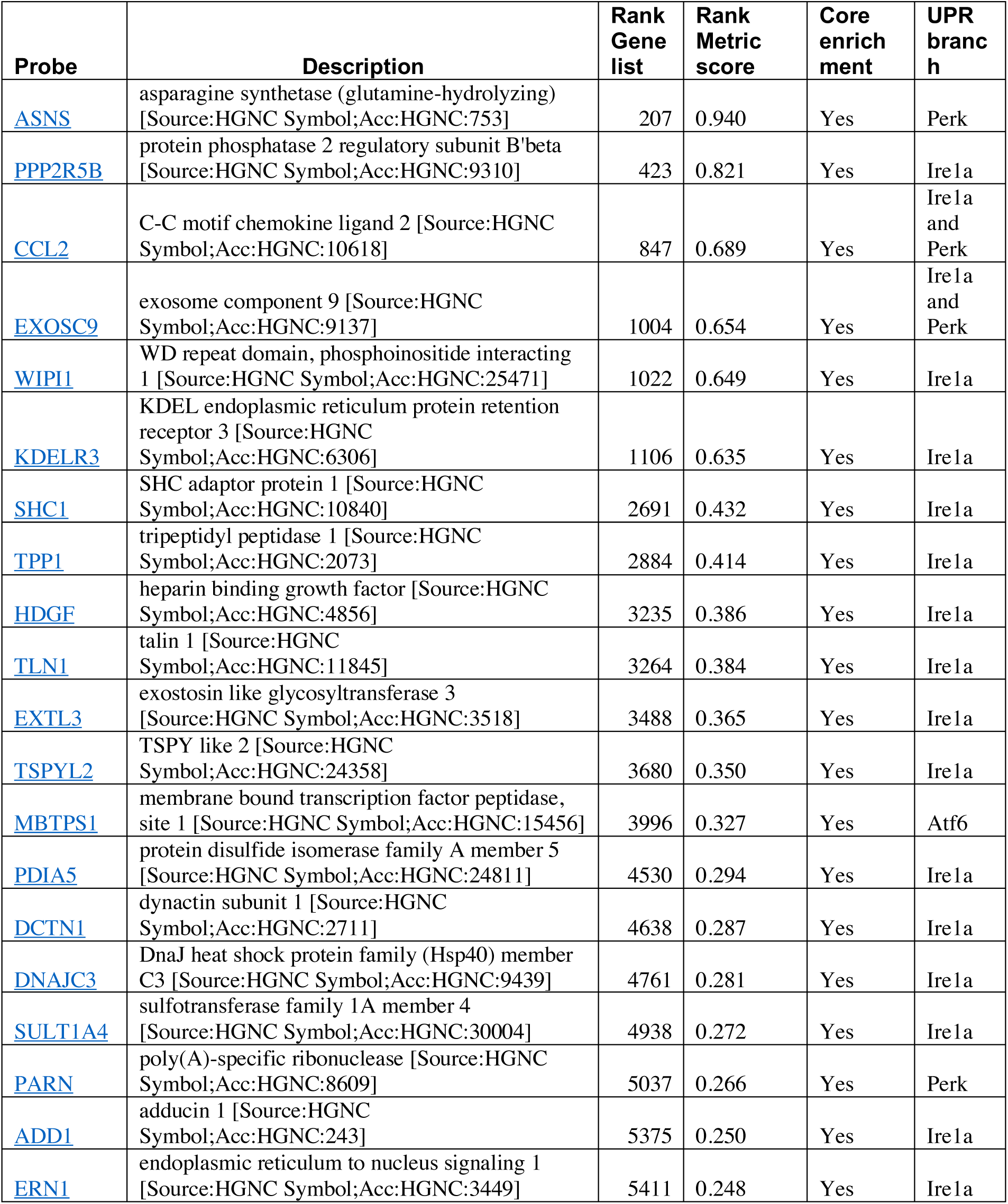
Genes the contributed to the core-enrichment of the GSEA.

### Tumor cells secrete factors that induce ER-stress in hepatic stellate cells

Hepatic stellate cell-lines (LX2) and HCC-cell lines (HepG2 and Huh7) were grown in different compartments using a transwell assay. This confirmed that tumor cells secrete factors that increase mRNA-expression of CHOP (Figure 2A), spliced XBP1 (Figure 2B and D) and BiP (Figure 2C), as well as protein expression of p-IRE1α (Figure 2E) in hepatic stellate cells co-cultured with tumor cells, indicating the presence of ER-stress. This also led to their activation, as measured by mRNA-expression of αSMA (Figure 2F) and collagen (Figure 2G) in LX2-cells grown with HepG2 or Huh7 cells in a transwell assay. The mRNA-expression of αSMA and collagen was restored to baseline levels when 4μ8C was added to the transwell co-cultures.

**Fig. 2.**
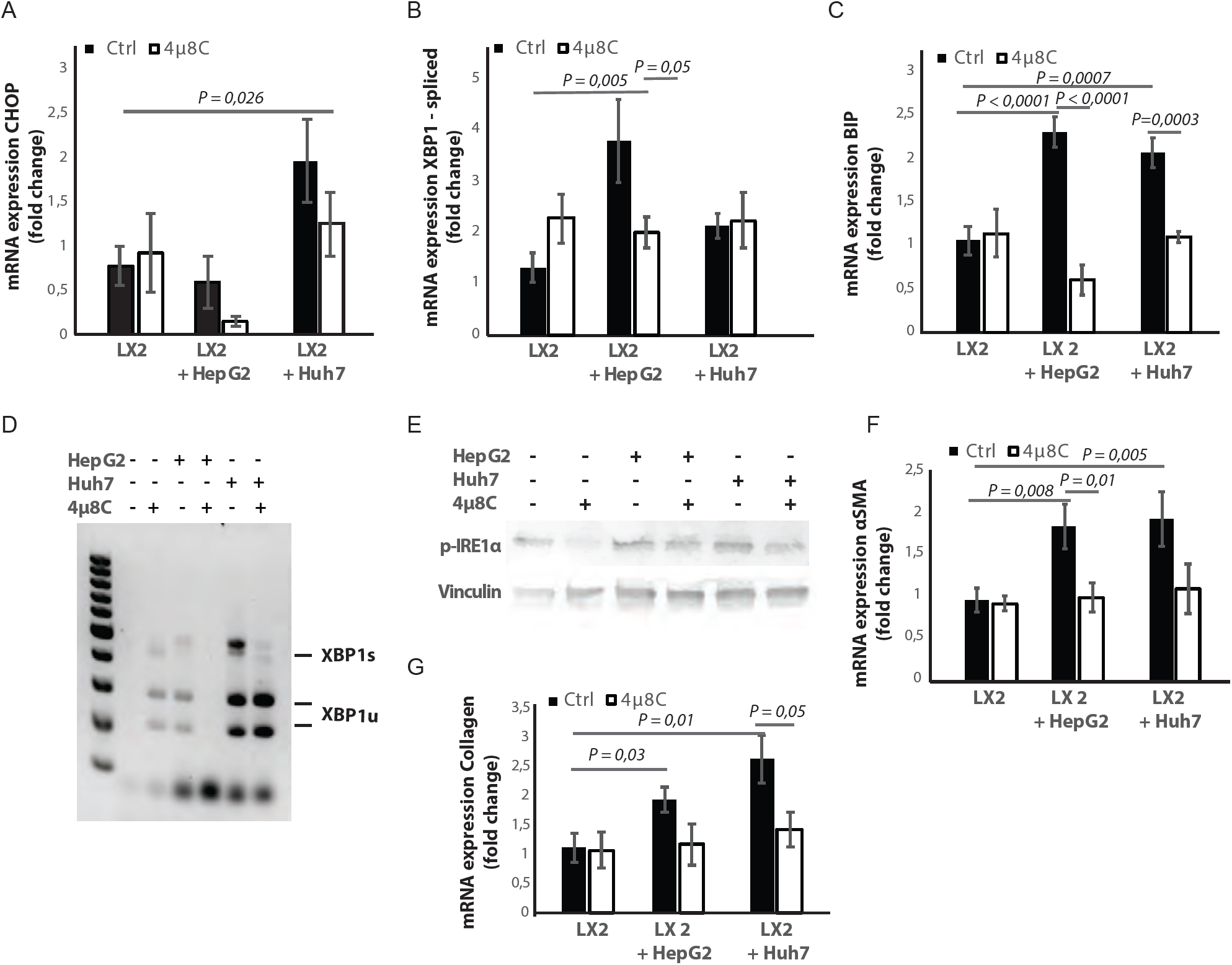
Tumor cells secrete factors that induce ER-stress in stellate cells, which contributes to their activation. ***(A)*** mRNA-expression of ER-stress markers CHOP, ***(B)*** spliced XBP1, (***C)*** BiP in stellate cells (LX2) co-cultured with cancer cells (HepG2 or Huh7) and treated with 4μ8C or control. **(D)** Detection of spliced (XBP1s) and unspliced XBP1 (XBP1u) visualized by digestion of XBP1u by *Pst-I*. (***E)*** protein expression of p-IRE1α and vinculin in stellate cells (LX2) co-cultured with cancer cells (HepG2 or Huh7) in transwell assays and treated with 4μ8C or control. (***F)*** mRNA-expression of stellate cell activation markers αSMA and (***G)*** collagen in LX2-cells co-cultured with HepG2 or Huh7-cells and treated with or without 4μ8C. P-values were calculated via the Student’s T-test with 10 biological replicates per group.

De-cellularised human liver 3D-scaffolds were engrafted with hepatic stellate cells (LX2) and tumor cells (HepG2). Sirius red staining and H&E staining confirmed that that LX2-cells and HepG2-cells successfully engrafted the collagen-rich matrix of the decellularized human liver scaffolds (Figure 3A). Engrafting both LX2-stellate cells and HepG2-cancer cells led to a significant increase of mRNA-expression of collagen, BiP and spliced XBP1 (Figure 3B) compared to scaffolds that were only engrafted with LX2-cells. Adding 4μ8C significantly decreased mRNA expression of collagen and BiP-mRNA-expression in the LX2 and HepG2 co-cultured scaffolds (Figure 3B).

**Fig. 3.**
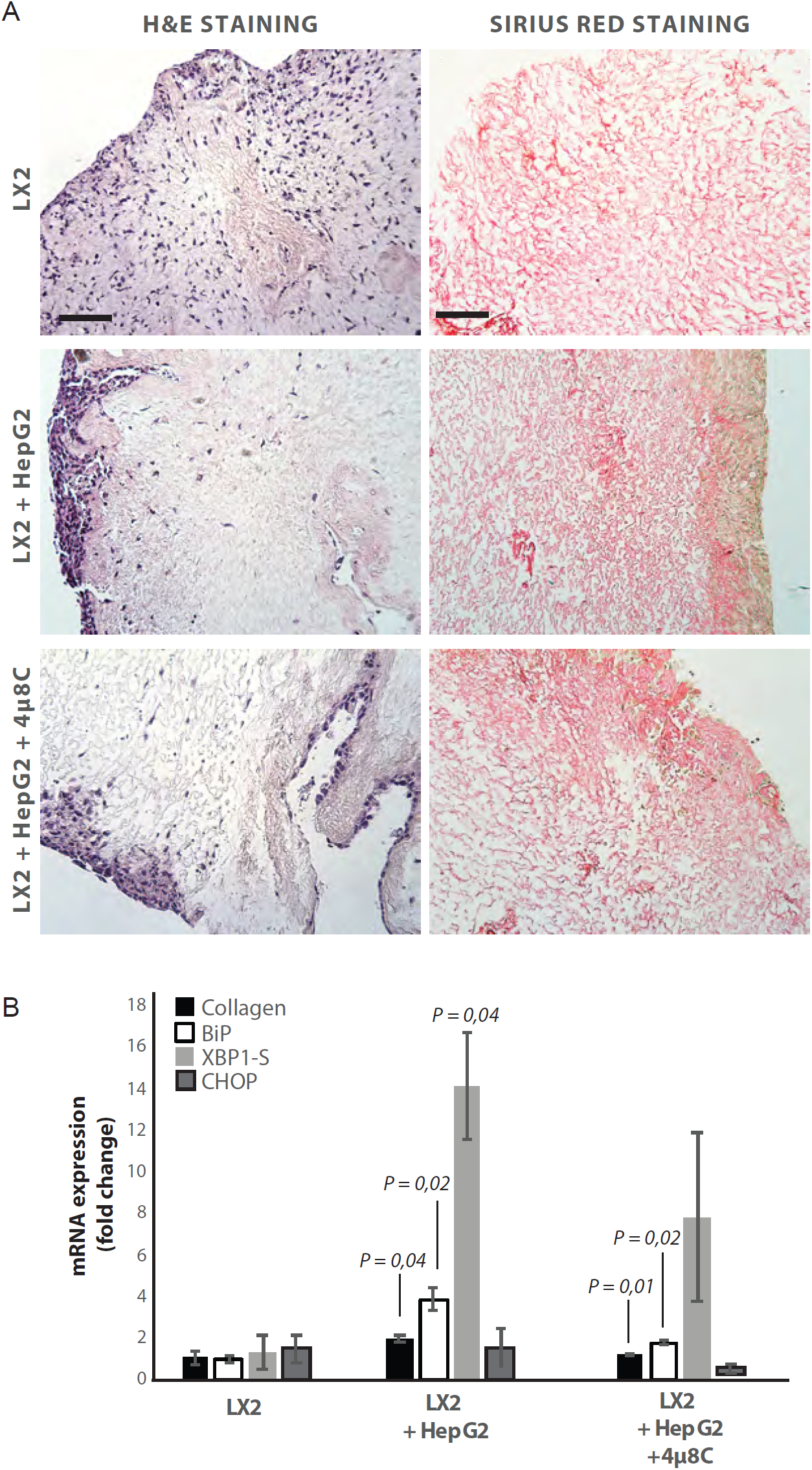
Inhibiting IRE1α decreases stellate cell activation in human liver 3D scaffolds engrafted with stellate cells and tumor cells. ***(A)*** Representative images of H&E and Sirius red stained slides of decellularized human liver scaffolds engrafted with LX2 stellate cells and HepG2-tumor cells treated with 4μ8C or control. ***(B)*** mRNA-expression of the stellate cells activation marker collagen and ER-stress markers BiP, spliced XBP-1 (XBP1-S) and CHOP in liver scaffolds engrafted with stellate cells (LX2) and cancer cells (HepG2), treated with 4μ8C or control. P-values were calculated via the Student’s T-test from 3 biological replicates per group, scale bars = 100μm.

Tumor cells are important sources of TGFβ, which is a known activator of stellate cells. Surprisingly, measuring TGFβ in mono-cultures lead to undetectable levels of TGFβ in Huh7-cells and low-levels in HepG2-cells (Supplementary figure 3A). These levels increased when LX2-cells were added to the co-cultures (Supplementary figure 3A). Engrafting both LX2-stellate cells and HepG2-cancer cells in the human liver scaffolds, slightly increased TGFβ-levels in the medium compared to scaffolds engrafted by only one cell type, but overall no significant differences were seen (Supplementary figure 3B). It is important to note that the baseline TGFβ-levels were markedly higher in the mono-cultured scaffolds, compared to the levels measured in cells grown in a standard 2D *in vitro* set-up (Supplementary figure 3A). Blocking TGFβ-receptor signaling with SB-431541 significantly reduced mRNA-expression of ER-stress markers CHOP (Supplementary figure 3C), spliced XBP1 (Supplementary figure 3D-E) and BiP (Supplementary figure 3F) in stellate cells co-cultured with tumor cells using transwells. Adding a TGFβ-receptor-inhibitor to stellate cell – tumor cell co-cultures also reduced stellate cell activation, as measured by mRNA-expression of αSMA (Supplementary figure 3G) and collagen (Supplementary figure 3H). This indicates that TGFβ-secretion by tumor cells could be responsible for activating stellate cells and for inducing the UPR.

### Pharmacological inhibition of IRE1α decreases tumor cell proliferation in stellate cell – tumor cell co-cultures

In transwell co-culturing assays, we found that co-culturing HepG2 or Huh7-tumor cells with LX2-stellate cells significantly increased PCNA-mRNA-expression in HepG2 and Huh7-tumor cell lines (Figure 4A). Adding 4μ8C significantly decreased mRNA-expression of PCNA in Huh7-cells grown in a transwell co-culture with LX2-cells, while not affecting PCNA-expression in tumor cell mono-cultures (Figure 4A). PCNA-levels in HepG2-LX2 transwell co-cultures were slightly decreased, but this was not significant. Proliferation was measured 24h after exposure to 4μ8C in tumor cells (HepG2 and Huh7) grown as mono-cultures and in co-culture with LX2-stellate cells. While 4μ8C induced a significant increase in proliferation of HepG2-monocultures, no difference was seen in LX2-monocultures and a significant decrease was seen in the HepG2-LX2 co-cultures (Figure 4B). In the Huh7 tumor cell line, 4μ8C significantly decreased cell number compared to untreated controls and a similar reduction was seen in the Huh7-LX2 co-cultures (Figure 4C). Immunohistochemical staining with antibodies against Epcam and Ki67 show that the effect on proliferation is mainly localized in the tumor cell population of these co-cultures (Figure 4D).

**Fig. 4.**
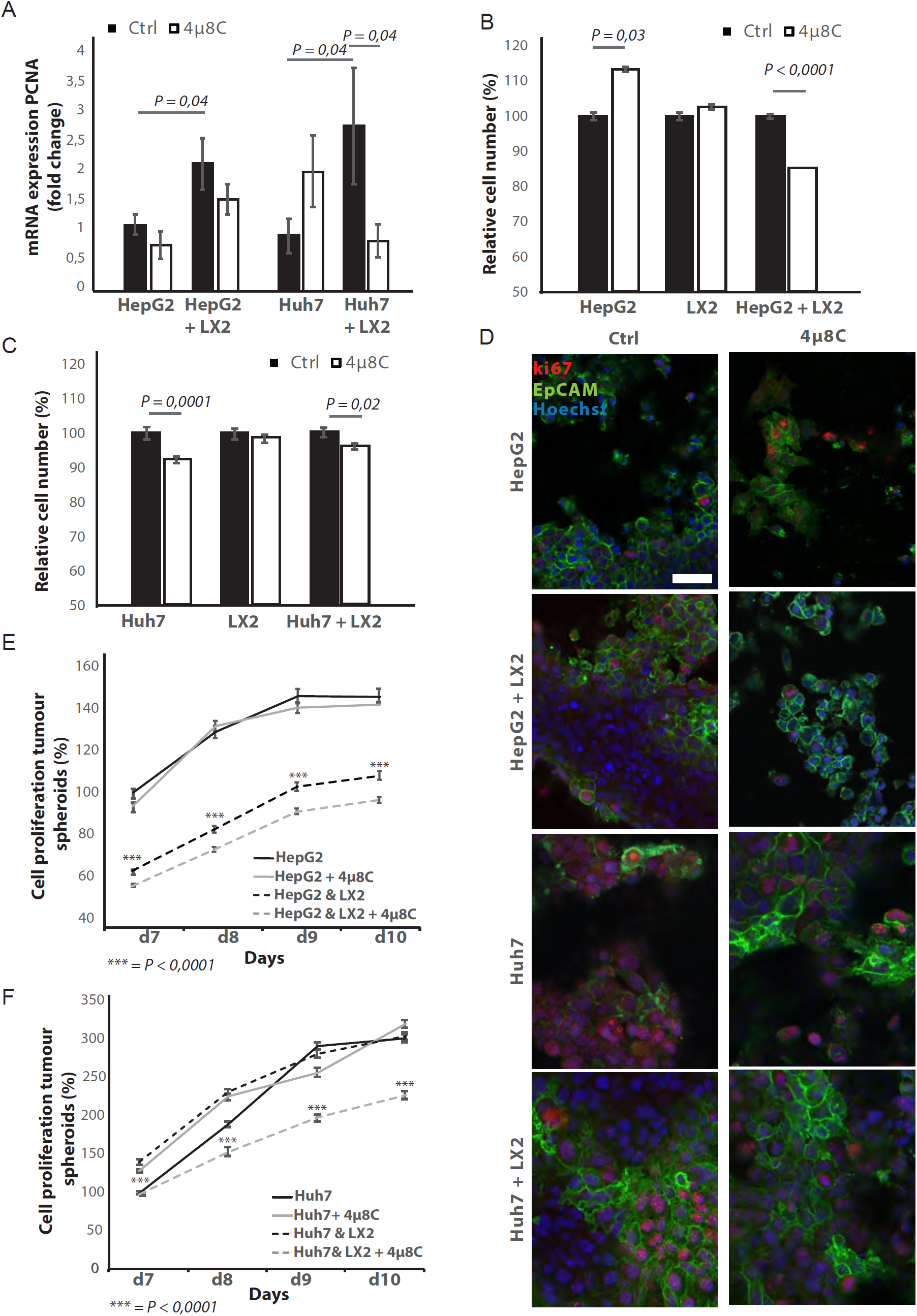
Inhibition of IRE1α decreases tumor cell proliferation. ***(A)*** PCNA mRNA-expression of HepG2 or Huh7-cells grown with LX2-cells in transwell inserts and treated with the IRE1α-inhibitor 4μ8C or control. ***(B)*** Relative cell number of LX2 and HepG2 or ***(C)*** LX2 and Huh7-cells treated with 4μ8C or control. ***(D)*** Representative images of tumor cells (HepG2 or Huh7) and LX2-stellate cells stained with antibodies against the HCC-marker Epcam and the proliferation marker ki67. ***(E)*** Cell proliferation of HepG2 or HepG2+LX2 spheroids and ***(F)*** Huh7 or Huh7+LX2 spheroids treated with 4μ8C or control. P-values were calculated via the Student’s T-test from 9 biological replicates per group, scale bars = 50μm.

3D-spheroids were generated using tumor cells alone (HepG2 or Huh7) or in combination with LX2-cells. While the HepG2-spheroids experienced a lower proliferation rate when co-cultured with LX2 stellate cells (Figure 4E), there was no difference in proliferation between spheroid-monocultures and spheroid-co-cultures in the Huh7-cells (Figure 4F). Treatment with 4μ8C significantly decreased proliferation of the tumor spheroids consisting of tumor cells (Huh7 or HepG2) and stellate cells (LX2), while tumor spheroid monocultures were not affected by 4μ8C. Similarly, PCNA-mRNA-expression significantly increased in human liver scaffolds engrafted with HepG2 and LX2-cells, compared to those engrafted with only tumor cells (Figure 5A). Treatment with 4μ8C significantly decreased PCNA-mRNA-expression in the LX2+HepG2 liver scaffolds, whilst not affecting those engrafted with only tumor cells. This further confirms our hypothesis that 4μ8C affects tumor cell proliferation indirectly, namely by blocking the activation of stellate cells and thus impairing the interaction between tumor and stroma.

**Fig. 5.**
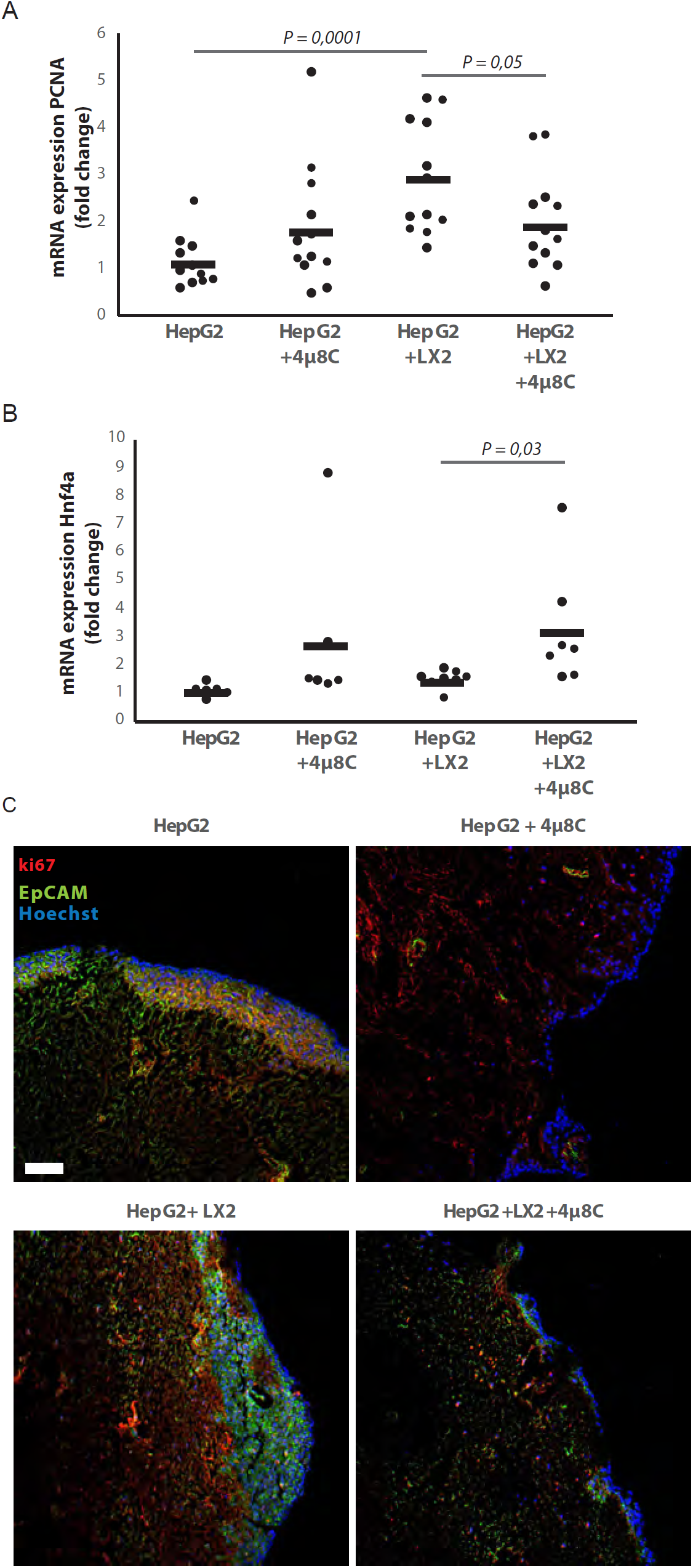
Inhibition of IRE1α decreases cell proliferation and improves liver function in human liver scaffolds engrafted with stellate cells and tumor cells. ***(A)*** PCNA and ***(B)*** Hnf4a expression of human liver scaffolds engrafted with HepG2-tumor cells and LX2-stellate cells, treated with 4μ8C or control. ***(C)*** Representative images of tumor cells (HepG2) and LX2-stellate cells stained with antibodies against the HCC-marker Epcam and the proliferation marker ki67. P-values were calculated via the Student’s T-test, scale bars = 100μm.

We measured the mRNA-expression of hepatocyte-nuclear-factor-4-alpha (Hnf4-α), which is a liver function marker that is correlated to a favorable outcome for HCC-patients (42). While co-engraftment of LX2 and HepG2-cells in the liver scaffolds only lead to a marginal increase of Hnf4-α, treatment with 4μ8C significantly increased Hnf4-α-mRNA-expression, thus suggesting an overall improvement of liver function and possibly improved prognosis (Figure 5B). Immunohistochemical staining of Epcam and ki67, showed that the HCC-cells have successfully engrafted the entire surface of the scaffolds and that 4μ8C decreases proliferation (Figure 5C).

### Pharmacological inhibition of IRE1α decreases tumor cell migration in stellate cell – tumor cell co-cultures

Co-culturing HepG2 and Huh7-tumor cells with LX2-cells in the transwell assays significantly increased mRNA-expression of the pro-metastatic marker MMP9 in HepG2-cells (Figure 6A) and MMP1 in HepG2 and Huh7-cells (Figure 6B). Adding 4μ8C significantly decreased the mRNA-expression of MMP1 in HepG2+LX2 and Huh7+LX2 transwell co-cultures, while a non-significant decrease of MMP9 mRNA-expression was seen in Huh7+LX2 transwell co-cultures. To assess whether this reduction in mRNA-expression of pro-metastatic markers has a functional effect on cell migration, a scratch wound assay was performed on confluent monolayers of mono-cultures (HepG2 or LX2) or tumor cell (HepG2) – stellate cell (LX2) co-cultures (Figure 6C). To visualize closing of the scratch wound by each individual cell type, cells were fluorescently labeled using CellTracker Green (tumor cells) or CellTracker Red (LX2 stellate cells) (Figure 6D). Tumor-stellate cell co-cultures were the most efficient to close the scratch wound (Figure 6E). This was significantly inhibited when co-cultures were treated with 4μ8C. We also observed a direct effect of 4μ8C on LX2 and HepG2-migration, since treatment with 4μ8C lead to a significant reduction in wound closure after 24h, compared to untreated controls. It is important to note that traditional scratch wound assays cannot distinguish between proliferation and migration (43). To overcome this limitation (44), we counted the individual number of cells in the middle of the wound area (Figure 6F and G). No significant difference was seen between HepG2 or LX2-cells within the wound area of HepG2-LX2 co-cultures after 24 hours (Figure9F). However, 4μ8C-treatment significantly decreased migration of HepG2-cells and LX2-cells inside the scratch wound in co-cultures, while not affecting mono-cultures (Figure 6G).

**Fig. 6.**
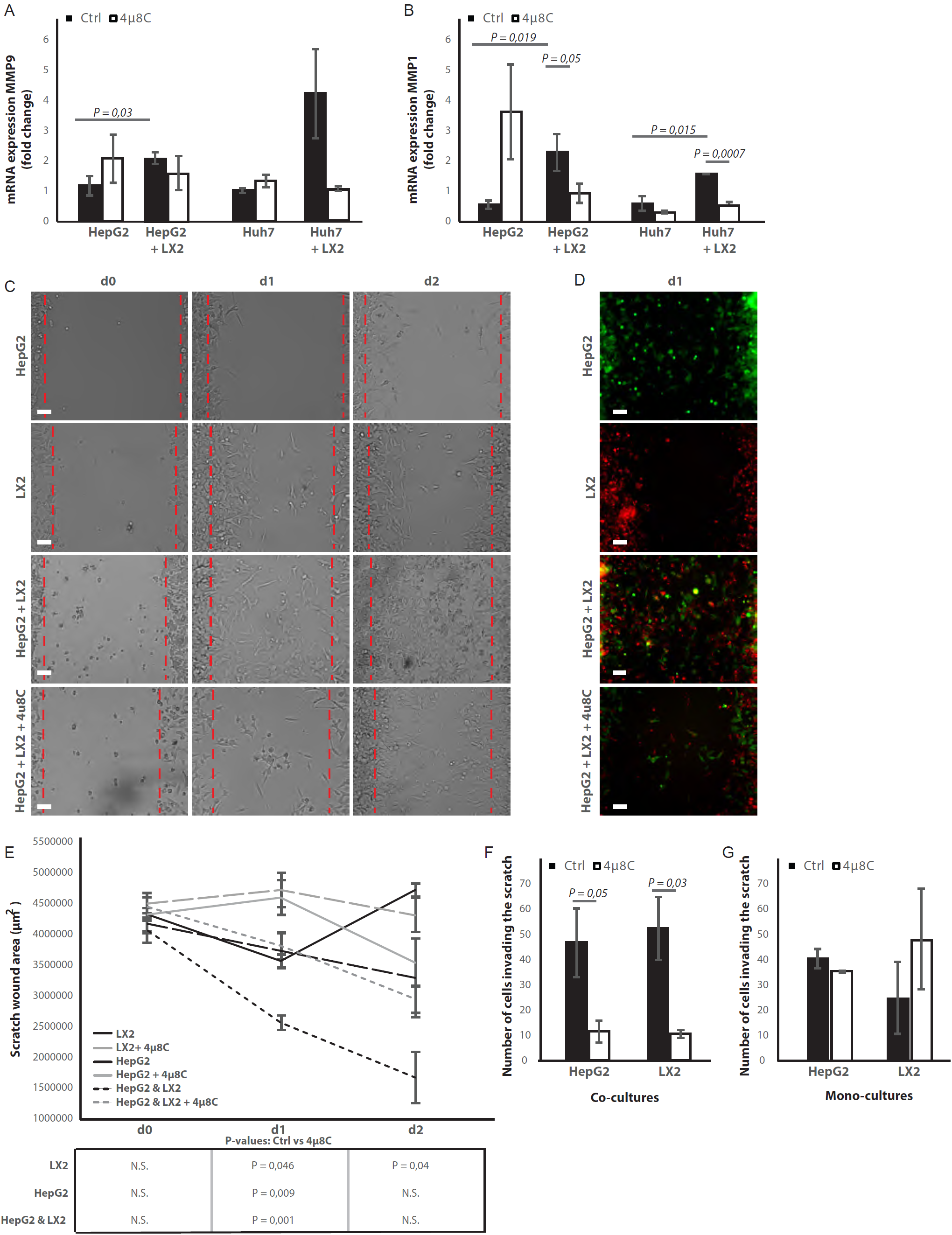
Inhibition of IRE1α decreases cell migration. ***(A)*** mRNA-expression of pro-metastatic markers MMP9 and ***(B)*** MMP1 in HepG2 and Huh7-cells co-cultured with LX2-cells and treated with 4μ8C or control. ***(C)*** Scratch wound on HepG2-cells and LX2-cells treated with 4μ8C or control. ***(D)*** Images of Cell Tracker stained HepG2-cells (Green) and LX2-cells (Red) invading the scratch area. ***(E)*** Quantification of wound size in HepG2-cells and LX2-cells treated with 4μ8C or control. ***(F)*** Number of HepG2-cells and LX2-cells invading the scratch wound after 24h in co-cultures and ***(G)*** mono-cultures. P-values were calculated via the Student’s T-test from 10 biological replicates per group (panel A and B) or 6 biological replicates per group (panel E-G), scale bars = 120μm.

Metastasis is usually a result of directed migration and chemotaxis toward physical and biochemical gradients within the tumor stroma (45). We used a microfluidic-based device for studying cell migration towards a stable gradient of chemotactic factors, such as FBS. 4μ8C significantly decreased total migration (Supplementary figure 4A-C) and directional migration towards FBS (Supplementary figure 4B and D) of HepG2-cells co-cultured with LX2-cells. Similarly, inhibition of ER-stress with 4μ8C significantly decreased total migration (Supplementary figure 4E and G) and directional migration towards FBS (Supplementary figure 4F and H) of LX2-cells co-cultured with HepG2-cells. Overall, these data suggest that stellate cells increase proliferation and pro-metastatic potential of tumor cells and blocking the IRE1α-RNase activity decreases tumor cell proliferation and migration.

### Silencing of IRE1α in stellate cells decreases tumor cell proliferation and migration in co-cultures

To investigate whether the effect of blocking IRE1α is due to a direct effect on the tumor cells or because of an indirect effect via stellate cells, we transfected the stellate-line LX2 with an IRE1α-siRNA prior to co-culturing them in a transwell assay with HepG2-cells. Transfection efficiency was determined via qPCR and showed a 50% reduction in the IRE1α-mRNA-expression (Figure 7A) compared to non-transfected (Ctrl) or mock-transfected (Scr) controls. In the transwell co-culturing assay, we found that silencing IRE1α in the LX2-cells significantly decreased PCNA-mRNA-expression in HepG2-cells (Figure 7B). Silencing IRE1α in the LX2-cells lead to a significant reduction of proliferation in LX2-HepG2 co-cultures (Figure 7C) and LX2-HepG2 spheroids (Figure 7D). Immunocytochemical staining with αSMA-antibodies (Figure 7E), confirmed a significant reduction of αSMA after si-IRE1α-transfection of LX2-stellate cells in HepG2-LX2 spheroid co-cultures (Figure 7F). A scratch wound assay on HepG2-LX2 co-cultures verified that silencing of IRE1α in LX2-cells significantly reduced wound closure compared to non-transfected and mock-transfected stellate cells (Figure 7G - H). Overall, these data confirm that blocking the IRE1α-pathway in hepatic stellate cells decreases proliferation and pro-metastatic potential of tumor cells.

**Fig. 7.**
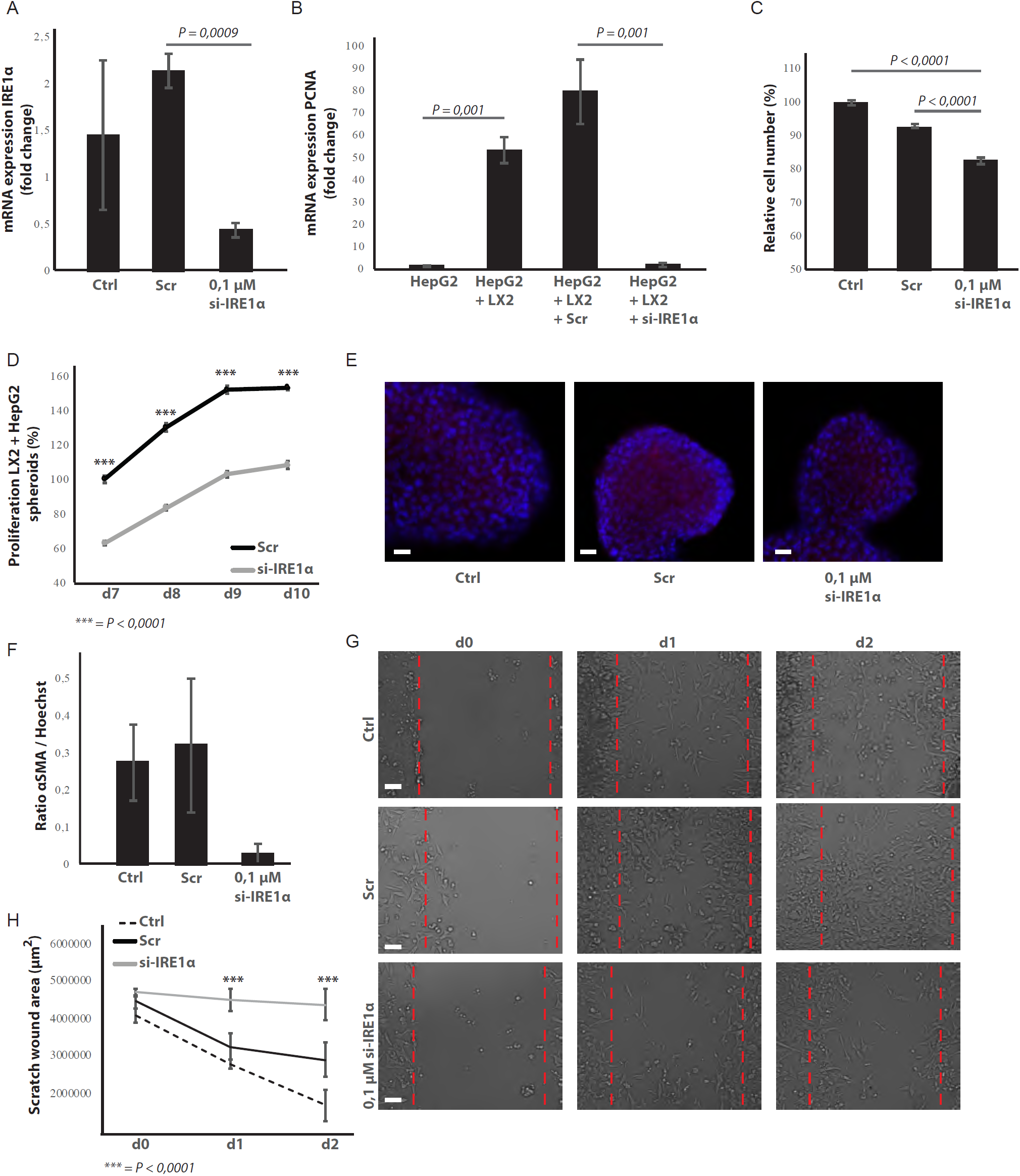
Silencing IRE1α in LX2-cells mimics 4μ8C. ***(A)*** IRE1α-mRNA-expression of LX2-cells transfected with IRE1α-siRNA (si-IRE1α), mock-transfected (Scr) or untransfected (Ctrl). ***(B)*** PCNA-mRNA-expression of HepG2-cells co-cultured with IRE1α-silenced LX2-cells or controls ***(C)***. Relative cell numbers in co-cultures of HepG2-cells and IRE1α-silenced LX2-cells or controls. ***(D)*** Proliferation of spheroids of HepG2-cells and IRE1α-silenced LX2-cells or controls ***(E)*** Images and ***(F)*** quantification of αSMA-stained spheroids with HepG2-cells and IRE1α-silenced LX2-cells or controls. ***(G)*** Images and ***(H)*** quantification of scratch wound of HepG2-cells co-cultured with IRE1α-silenced LX2-cells or controls. P-values were calculated via the Student’s T-test from 3 biological replicates per group, scale bars = 50μm (E) or 120μm (G).

## DISCUSSION

There is increasing evidence that ER-stress and activation of the UPR play an essential role during hepatic inflammation and chronic liver disease. We have previously shown that inhibition of IRE1α prevents stellate cell activation and reduces liver cirrhosis *in vivo* (25). In this report, we further define a role of ER-stress and the UPR in the interaction between tumor cells and hepatic stellate cells. We also show that IRE1α could form a valuable therapeutic target to slow down the progression of hepatocellular carcinoma.

Activated stellate cells play an important role in promoting tumorigenesis and tumors are known to secrete cytokines such as TGFβ, which induce myofibroblast activation and creates an environment that sustains tumor growth (46). Since over 80% of HCC arises in a setting of chronic inflammation associated with liver fibrosis, targeting the fibrotic tumor micro-environment is often proposed as a valuable therapeutic strategy for HCC-patients (2). We and others have shown that ER-stress plays an important role in stellate cell activation and contributes to the progression of liver fibrosis (25, 47-49). The mechanisms by which the UPR promotes stellate cell activation have been attributed to regulating the expression of c-MYB (25), increasing the expression of SMAD-proteins (47) and/or by triggering autophagy (49).

In our study, we show that ER-stress plays an important role in stellate cell – tumor cell interactions and that pharmacological inhibition of IRE1α-endoribonuclease activity slows down the progression of HCC *in vivo*. We demonstrate that tumor cells induce ER-stress in hepatic stellate cells, thereby contributing to their activation and creating an environment that is supportive for tumor growth and metastasis. Activated stellate cells are known to enhance migration and proliferation of tumor cells *in vitro (8)* and *in vivo* (50), possibly by producing extracellular matrix proteins and by producing growth factors. Extracellular matrix proteins such as collagen can act as a scaffold for tumor cell migration (51), alter the expression of MMP’s (8) and induce epithelial-mesenchymal transition (52). Activated stellate cells are also an important source of hepatocyte growth factor, which promotes proliferation, cell invasion and epithelial-mesenchymal transition via the c-MET signaling pathway (53). Interestingly, blocking ER-stress in the stellate cell population reduced tumor-induced activation towards myofibroblasts, which then decreases proliferation and migration of tumor-cells in co-cultures. This suggests that targeting the microenvironment using an ER-stress inhibitor could be a promising strategy for patients with HCC.

The UPR has been described as an essential hallmark of HCC (54), although its role within tumorigenesis remains controversial (18). While a mild to moderate level to ER-stress leads to activation of the UPR and enables cancer cells to survive and adapt to adverse environmental conditions, the occurrence of severe or sustained ER-stress leads to apoptosis. Both ER-stress inhibitors as ER-stress inducers have therefore been shown to act as potential anti-cancer therapies (55). A recent study by Wu *et al*, demonstrated that IRE1α promotes progression of HCC and that hepatocyte specific ablation of IRE1α results in a decreased tumorigenesis (56). In contrast to their study, we found a greater upregulation of actors of the IRE1α-branch within the stroma than in the tumor itself and identified that expression of ER-stress markers was mainly localized within the stellate cell population. An important difference between both studies is the mouse model that is used. While Wu *et al* used a single injection of DEN, we performed weekly injections, causing tumors to occur in a background of fibrosis, similar to what is seen in patients (26). Our *in vitro* studies with mono-cultures confirm that 4μ8C also has a direct effect on proliferation and migration of HCC cells – similar to the findings of Wu *et al* - and the response seems to depend on the tumor cell line. Adding 4μ8C to HepG2-cells significantly increased proliferation, while a significant decrease was seen in the Huh7-cells. This difference in response could be due IRE1α’s function as a key cell fate regulator. On the one hand it can induce mechanisms that restore protein homeostasis and promote cytoprotection, while on the other hand IRE1α also activates apoptotic signaling pathways. How and when IRE1α exerts its cytoprotective or its pro-apoptotic function remains largely unknown. The duration and severity of ER-stress seems to be a major contributor to the switch towards apoptosis, possibly by inducing changes in the conformational structure of IRE1α (57). The threshold at which cells experience a severe and prolonged ER-stress that would induce apoptosis could differ between different cell lines, depending on the translational capacity of the cells (e.g. ER-size, number of chaperones and the amount of degradation machinery) and the intrinsic sources that cause ER-stress (58). A study of Li *et al*, has specifically looked at how IRE1α regulates cell growth and apoptosis in HepG2-cells (59). Similar to our findings, they discovered that inhibiting IRE1α enhances cell proliferation, while over-expression of IRE1α increases the expression of polo-like kinase, which leads to apoptosis. Interestingly, polo-like kinases have divergent roles on HCC-cell growth depending on which cell line is used, which could explain the different response to 4μ8C in Huh7 and HepG2-cells (60). Studies on glioma cells show that IRE1α regulates invasion through MMP’s (61). In line with these results, we also detected a reduction of MMP1-mRNA expression after 4μ8C-treatment and observed a direct effect on wound closure in HepG2-cells. These results indicate that ER-stress could play a direct role in regulating tumor cell invasion, in addition to its indirect effect via stellate cells.

In conclusion, the aim of this study was to define the role of ER-stress in the cross-talk between hepatic stellate cells and tumor cells in liver cancer. We show that pharmacologic inhibition of the IRE1α-signaling pathway decreases tumor burden in a DEN-induced mouse model for HCC. Using several *in vitro* 2D and 3D co-culturing methods, we identified that tumor cells induce ER-stress in hepatic stellate cells and that this contributes to their activation. Blocking ER-stress in these hepatic stellate cells prevents their activation, which then decreases proliferation and migration of tumor cells.

## Abbreviations

αSMA: α-smooth muscle actin;
DEN: diethylnitrosamine;
DMEM: Dulbecco modified eagle medium;
ELISA: Enzyme-Linked immune Sorbent Assay,
ER: Endoplasmic reticulum;
FBS: fetal bovine serum;
HCC: Hepatocellular carcinoma,
H&E: Haematoxilin-eosin;
TBS: tris-buffer saline;
TGFβ: tumor growth factor β;
UPR: unfolded protein response;

## ACKNOWLEDGEMENTS

This research was funded through grants obtained from the Swedish Cancer Foundation (Cancerfonden, CAN2017/518 and CAN2013/1273), The Swedish children’s cancer foundation (Barncancerfonden), the Swedish society for medical research (SSMF, S17-0092), the O.E. och Edla Johanssons stiftelse. These funding sources were not involved in the study design; collection, analysis and interpretation of data; writing of the report; and in the decision to submit the article for publication. We would like to thank visiting students Kim Vanhollebeke and Justine Dobbelaere for their technical assistance; GradienTech for providing us with their CellDirector assays and Paul O’Callaghan for his valuable input on our project.

## Competing interest

The authors have no conflict of interest to report.

## Figure legends

**Supplementary figure 1. Activation of the unfolded protein response is mainly located in the stroma of mice with HCC. *(A)*** mRNA-expression of the ER-stress marker BiP in tumor and surrounding non-tumoral liver tissue in mice with DEN-induced HCC treated with or without the IRE1α-inhibitor 4μ8C or healthy mice. ***(B)*** Representative western blot image showing protein expression of BiP in liver tissue from mice with HCC treated with or without 4μ8C **(C)** Quantification of p-IRE1α staining on murine liver sections. ***(D)*** Co-staining of liver tissue with antibodies against αSMA and p-IRE1α. ***(E)*** Co-staining of liver tissue with antibodies against αSMA and BiP. P-values were calculated via the Student’s T-test, scale bars = 50μm.

**Supplementary figure 2. Activation of the unfolded protein response pathway is increased in patients with fibrotic HCC. *(A)*** Heat map showing gene-set enrichment analysis results from samples from fibrous HCC versus non-fibrous HCC. ***(C)*** Immunohistochemically stained liver biopsies from HCC-patients obtained from the human protein atlas, using antibodies against IRE1α-mediated actors of the unfolded protein response: WIPI1, SHC1, PPP2R5B and BiP. (D) Kaplan-Meier survival curves of HCC-patients with high or low expression of WIPI1, SHC1, PPP2R5B and BiP. P-values were calculated via a Log-Rank test.

**Supplementary Figure 3. Secretion of TGFβ by tumor cells activates stellate cells and induces ER-stress. *(A)*** concentration of TGFβ in medium from tumor cells (HepG2 or Huh7) grown in mono-culture or co-cultured with LX2-stellate cells, treated with 4μ8C or control. ***(B)*** concentration of TGFβ in medium from liver scaffolds engrafted with stellate cells ***(C)*** (LX2) and tumor cells (HepG2) treated with 4μ8C or control. mRNA-expression of the ER-stress markers CHOP, ***(D)*** spliced XBP1, ***(E)*** unspliced XBP1 and ***(F)*** BiP in hepatic stellate cells (LX2) grown as mono-culture or in co-cultures with the cancer cell lines HepG2 and Huh7 treated with the TGFβ receptor inhibitor SB-431541 or control. (***G)*** mRNA-expression of stellate cell activation markers αSMA and (***H)*** collagen in LX2-cells grown with HepG2 or Huh7-cells and treated with SB-431541 or control. P-values were calculated via the Student’s T-test from 7 biological replicates per group.

**Supplementary Figure 4. Inhibiting IRE1α decreases chemotaxis. *(A)*** migration plots of LX2-cells co-cultured with HepG2-cells exposed to an FBS-gradient (increasing towards the right) and treated with control or ***(B)*** 4μ8C ***(C)*** Quantification of total migration and ***(D)*** directional migration of LX2-cells (co-cultured with HepG2-cells) towards an FBS-gradient with or without 4μ8C. ***(E)*** Migration plots of HepG2-cells co-cultured with LX2-stellate cells and exposed to an FBS-gradient and treated with control or ***(F)*** 4μ8C. ***(G)*** Quantification of total migration and ***(H)*** directional migration of HepG2-cells (co-cultured with LX2-cells) towards an FBS-gradient with or without 4μ8C. P-values were calculated via the Student’s T-test from 3 biological replicates per group.

**Supplementary Table 1. Primer sequences**

**Supplementary methods 1: histology and immunohistochemistry**

## Notes

#### Summary of Updates

Minor changes in the text to fulfil the manuscript guidelines for full submission in eLife.

