## Supplementary figures and images for "Inhibiting endoplasmic reticulum stress decreases tumor burden in a mouse model for hepatocellular carcinoma"

### Supplementary figure 1

#### Supplementary figure 1

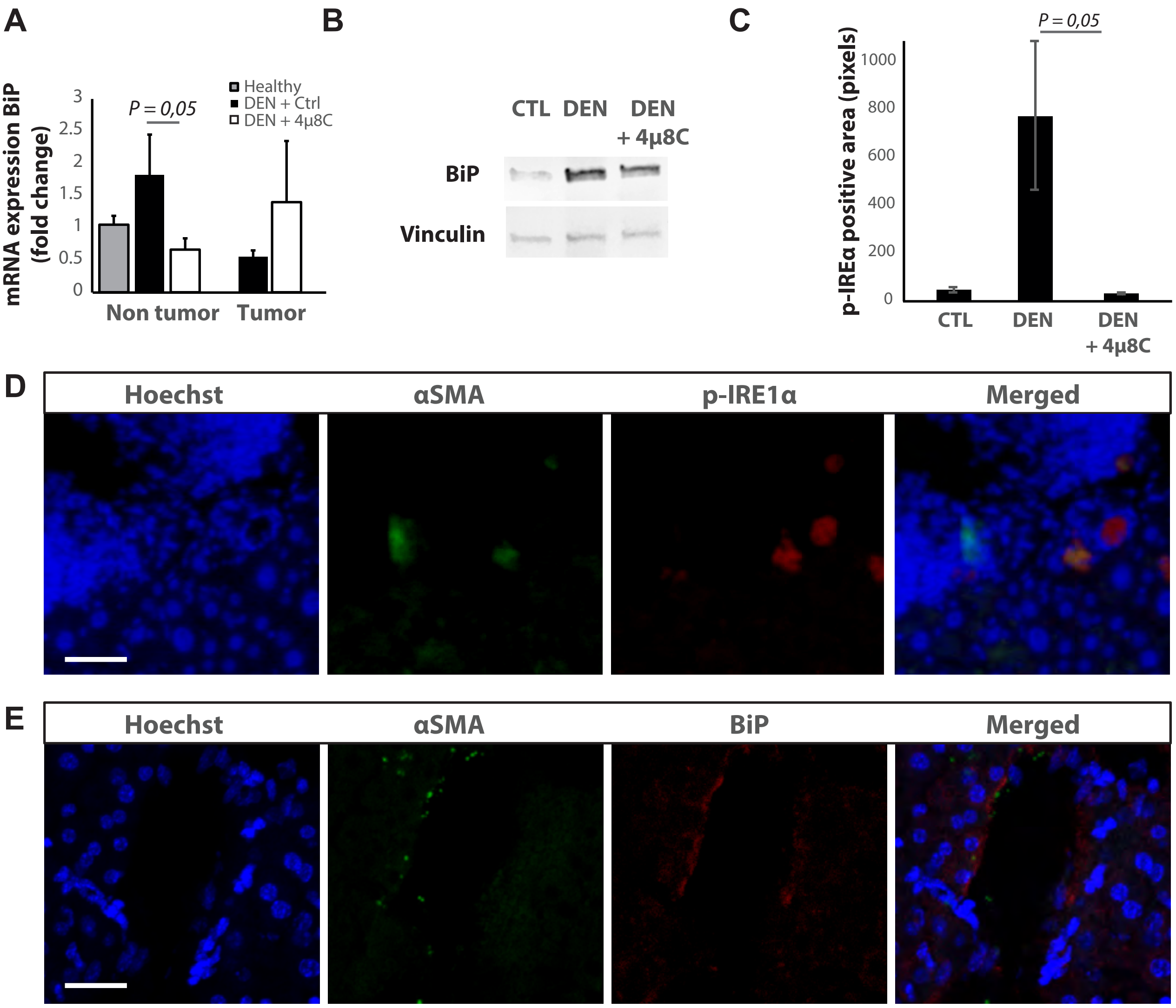

### Supplementary figure 2

##### Supplementary figure 2

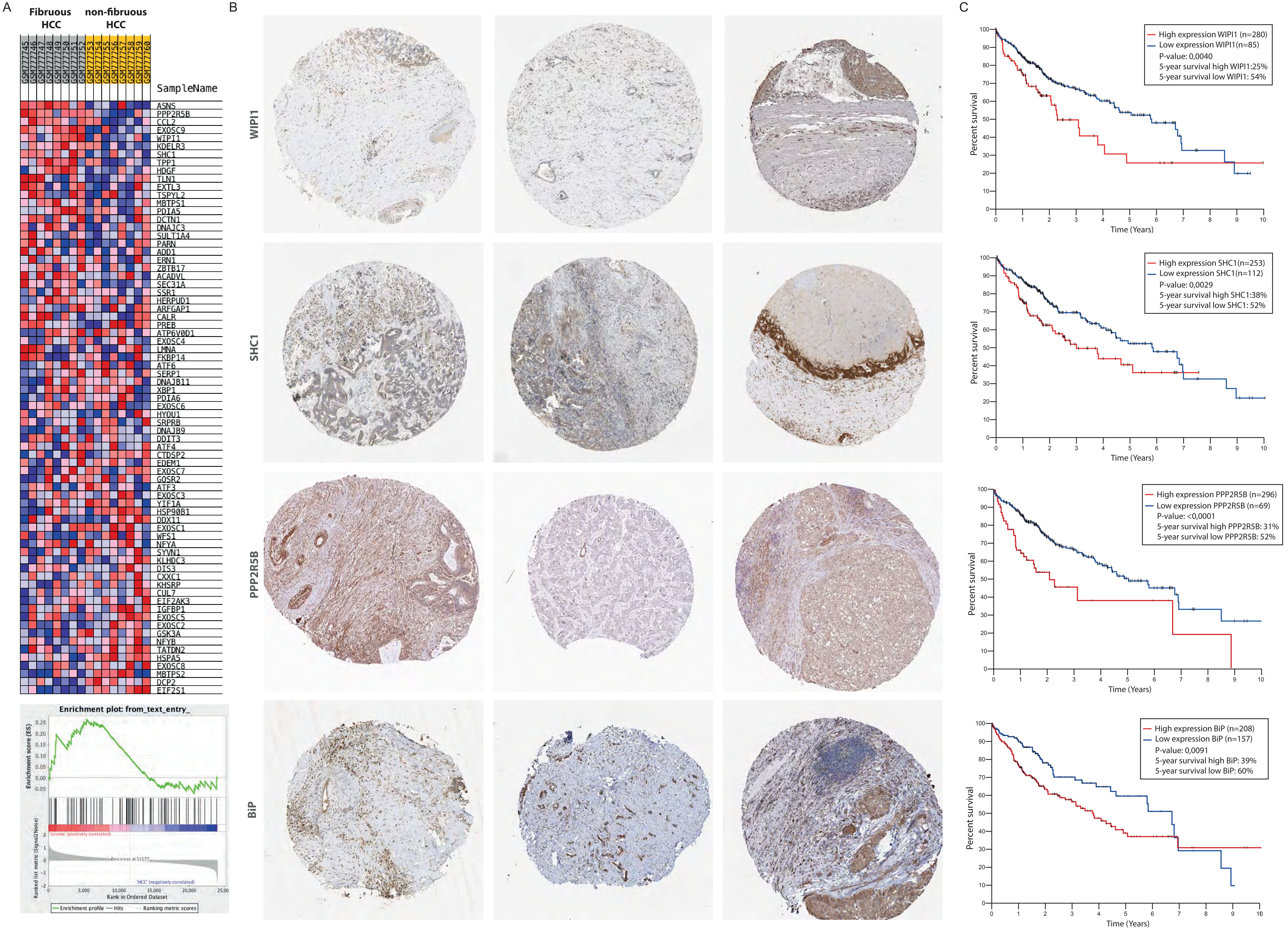

### Supplementary figure 3

#### Supplementary figure 3

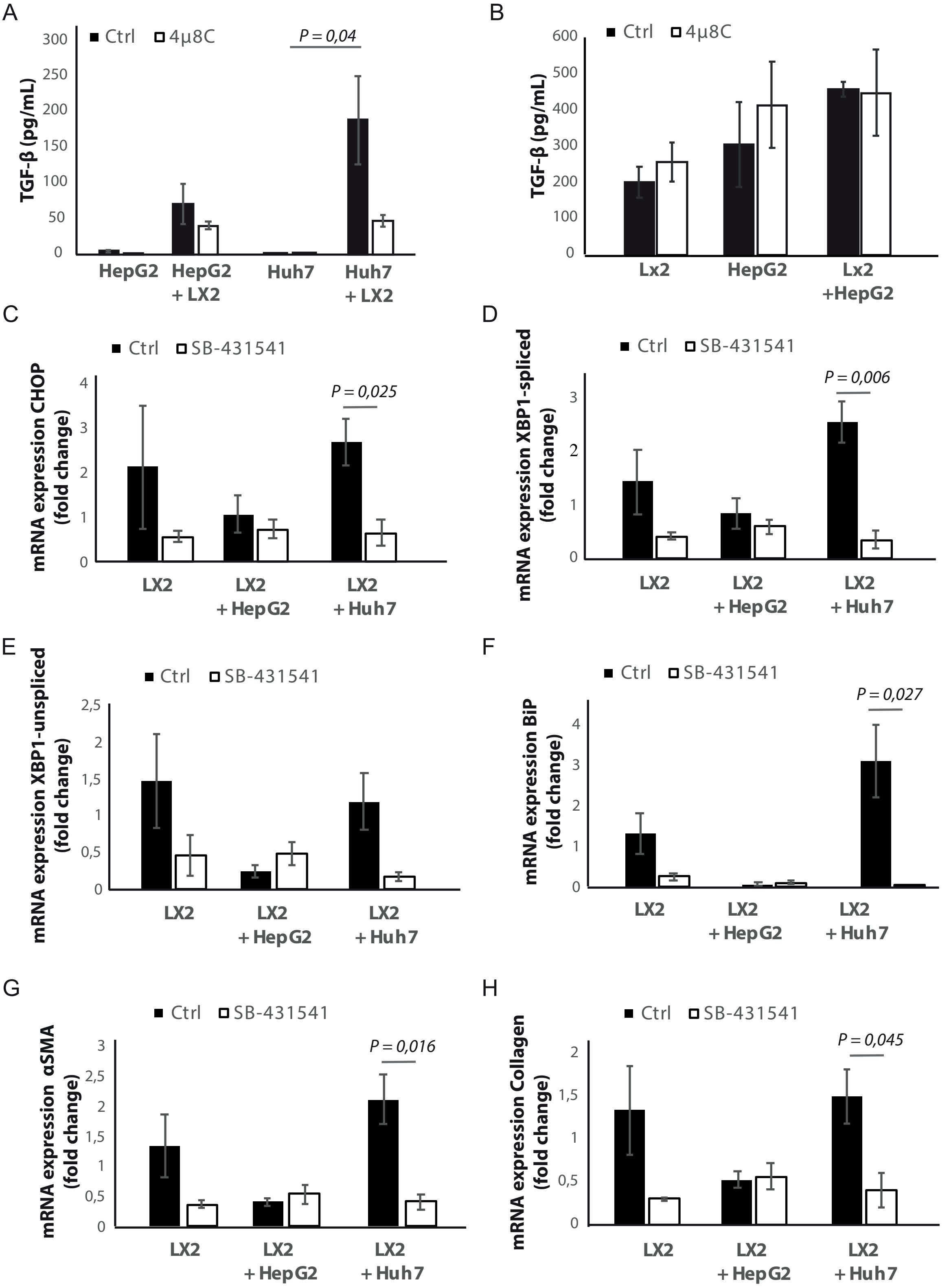

### Supplementary figure 4

#### Supplementary figure 4

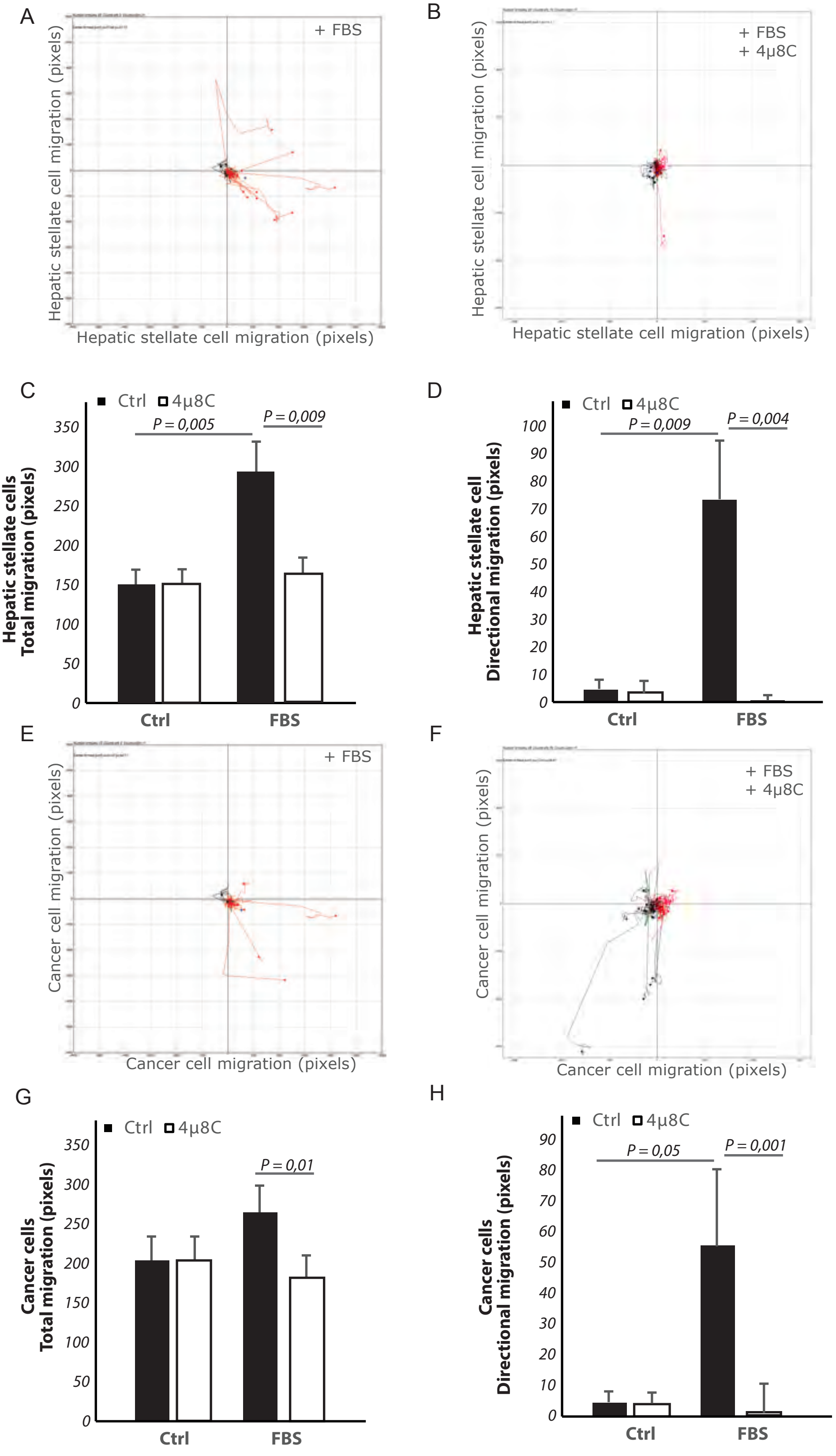
